# Deciphering the nanoscale architecture of presynaptic actin using a micropatterned presynapse-on-glass model

**DOI:** 10.1101/2024.09.05.611287

**Authors:** Sofia Tumminia, Louisa Mezache, Theresa Wiesner, Benoit Vianay, Manuel Théry, Marie-Jeanne Papandréou, Christophe Leterrier

## Abstract

Chemical synapses are fundamental units for the transmission of information throughout the nervous system. The cytoskeleton allows to build, maintain and transform both pre- and postsynaptic contacts, yet its organization and the role of its unique synaptic nanostructures are still poorly understood. Here we present a presynapse-on-glass model where presynaptic specializations are robustly induced along the axons of cultured neurons by micropatterned dots of neuroligin, allowing the controlled orientation and easy optical visualization of functional induced presynapses. We demonstrate the relevance and usefulness of this presynapse-on-glass model for the study of presynaptic actin architecture, showing that a majority of induced presynapses are enriched in actin, with this enrichment being correlated to higher synaptic cycling activity. We confirm our previous results on bead-induced presynapses by identifying the same distinct actin nanostructures within presynapses: corrals, rails and mesh. Furthermore, we leverage the controlled orientation of the presynapse-on-glass model, visualizing the arrangement of these actin structures relative to the active zone nanoclusters using multicolor 3D Single Molecule Localization Microscopy (SMLM), and relative to the sub-diffractive localization exocytic events using a correlative live-cell and SMLM approach.

## Introduction

Neuronal communication is a complex process involving coordinated interactions between cell-adhesion molecules, signaling complexes, and cytoskeletal elements, which are essential for precise signal transmission across chemical synapses (McAllister, 2007; Togashi et al., 2009). Noteworthy is the actin cytoskeleton, which is crucial in maintaining neuronal structure and shaping neurons throughout their development. It provides the essential framework that supports neuron morphology and enables dynamic changes necessary for growth and function during both early development and maturity (Luo, 2002; Coles and Bradke, 2015). Disruptions in these dynamics can impair neuronal polarization and axonal guidance, leading to ineffective neuronal communication and potentially contributing to neurodevelopmental and psychiatric disorders (Bernstein et al., 2010; Yan et al., 2016).

In addition to its structural role, actin dynamics are vital for neuronal function and synaptic transmission (Cingolani and Goda, 2008; Papandréou and Leterrier, 2018; Gentile et al., 2022). The actin cytoskeleton plays a significant role in pre-synaptic vesicle clustering, mobilization, and docking, which are essential for maintaining synaptic efficacy (Bernstein and Bamburg, 1989; Kim and Lisman, 1999; Doussau and Augustine, 2000). Actin is also important for endocytosis and the generation of synaptic vesicles at presynapses (Shupliakov et al., 2002; Watanabe et al., 2013; Ogunmowo et al., 2023). As regards the exocytic process, studies using short-term application of actin-perturbing agents have led to contradictory findings of reduced (Cole et al., 2000) or enhanced (Cole et al., 2000; Sankaranarayanan et al., 2003) vesicular release, suggesting that actin acts as a meshwork that gathers vesicles before exocytosis, or as a physical barrier within the presynaptic subdomains. These opposing claims underscore the need for further research to clarify its mechanisms (Li et al., 2018a).

A better understanding of actin’s roles at the presynapse requires knowledge of its structural architecture (Papandréou and Leterrier, 2018). Early ultrastructural studies using electron microscopy have provided significant insights into the distribution and organization of actin within synapses, revealing a dense meshwork of actin in dendritic spines (Matus et al., 1982), and shedding light on presynaptic components and actin interconnections within synaptic vesicular clusters (Landis et al., 1988; Hirokawa et al., 1989) and at the active zone periphery (Bloom et al., 2003). However, electron microscopy studies are limited by the difficulty distinguishing actin from other cytoskeletal elements, the complex sample preparation leading to potential damage, and the restrained molecular specificity inhibiting a comprehensive model of presynaptic architecture.

Fluorescence microscopy techniques have expanded our knowledge about the spatial relationship between presynaptic components due to their better molecular specificity, despite its diffraction-limited resolution being inadequate to resolve presynaptic cytoskeletal and molecular nanostructures. Fortunately, super-resolution microscopy approaches can overcome the diffraction limit and allow nanoscale imaging studies (Jacquemet et al., 2020; Werner et al., 2021). Among these, Single Molecule Localization Microscopy (SMLM) techniques such as Stochastic Optical Reconstruction Microscopy (STORM) and DNA-Point Accumulation in Nanoscale Topography (PAINT), which rely on stochastic blinking of single fluorophores followed by their precise localization, have refined our knowledge of presynaptic organization (Dani et al., 2010; Glebov et al., 2017; Carvalhais et al., 2021).

Despite those advances, the dense expression of actin at the postsynapse makes presynaptic actin imaging quite challenging in its natural context (Reshetniak and Rizzoli, 2019; Nosov et al., 2020). To overcome these obstacles, our team has recently used a model of bead-induced isolated presynapses (Burry, 1980; Lucido et al., 2009). By visualizing actin within isolated presynapses by SMLM, we characterized three distinct presynaptic actin nanostructures: a faint active zone mesh, linear actin rails found between the active zone and the reserve pools, as well as actin corrals that surround the presynaptic compartment (Bingham et al., 2023). However, presynapses formed on ∼5 µm spherical beads result in heterogeneous orientation and sizes of induced presynapses. This limits optical accessibility and unambiguous localization of the active zone, impeding the analysis of actin structure roles in dynamic processes like presynaptic exo-endocytosis. An alternative is to induce the formation of presynapses directly on the glass, using postsynaptic adhesion proteins such as neuroligin applied on the whole coverslip (Funahashi et al., 2018) or on micropatterns of SynCAM1 separated by cytophobic regions (Czöndör et al., 2013).

In this study, we designed and optimized a presynapse-on-glass model to orient presynaptic formation toward patterned dots displaying the extracellular domain of neuroligin-1 (nlgn) in addition to a continuous layer of polylysine and laminin. This allowed unrestricted growth of neuronal processes as well as robust induction of presynaptic specialization near the patterned nlgn. We demonstrate how this model recapitulates actin distribution and the nanoscale organization we observed in bead-induced and endogenously tagged presynapses (Bingham et al., 2023). Furthermore, presynapses-on-glass allow to refine the 3D visualization of actin nano-architecture at presynapses, and to combine the visualization of synaptic vesicle exocytosis in living neurons with the nanoscopy of presynaptic components to link actin nano-architecture to the precise location of vesicle exocytosis.

## Results

### Micropatterned neuroligin induces axonal presynaptic marker clustering and actin aggregation

To improve optical accessibility and homogeneity of presynaptic formation, we adapted a validated method of presynaptic induction, where coating the coverslip with the extracellular domain of a postsynaptic cell-adhesion molecule, neuroligin-1 fused to the N-terminal of human immunoglobulin-Fc region (nlgn) induces the formation of excitatory presynapses onto the glass (Funahashi et al., 2018). We used micro-contact printing (Théry, 2010) of a PDMS stamp with 1.8 µm diameter pillars to generate an array of ∼2 µm dots on the coverslip. To induce presynapses, the pillars were coated with nlgn together with fluorescent BSA for pattern visualization (nlgn dots). After printing, the coverslips were coated with a uniform layer of polylysine and laminin, so that axons and dendrites could grow freely and react when crossing printed dots. Rat hippocampal neurons were then cultured onto printed coverslips, and subsequently fixed and immunolabeled for presynaptic markers after 14-15 days in vitro (div) to examine presynaptic formation. Isolated axons containing at least three successive induced presynapses on neuroligin dots were classified as “patterned axons”. The formation of patterned axons was assessed (as the percentage of total length of isolated axons over the field of view) for two presynaptic proteins: the active zone scaffolding protein bassoon (Dieck et al., 1998) and the synaptic vesicle membrane protein synaptophysin (Jahn et al., 1985; Wiedenmann and Franke, 1985). In addition, active vesicular cycling was evaluated using feeding with a synaptotagmin antibody against an extracellular epitope that accumulates in presynapses through constitutive vesicular cycling (Kraszewski et al., 1995). Representative fluorescence images demonstrate aggregation of bassoon, synaptophysin and synaptotagmin uptake into axonal clusters near nlgn dots (Fig. 1A-F). All three markers yielded comparable occurrences of patterned axons along nlgn dots patterns (23 ± 3% with bassoon, 16 ± 3% with synaptophysin, and 23.3 ± 2% with synaptotagmin, Fig. 1G), while neurons cultured on patterns of dots containing a human Fc without fused neuroligin or outside of the micropatterned areas resulted in almost no patterned axons (1.5 ± 0.4%, Fig. S1).

**Figure 1:**
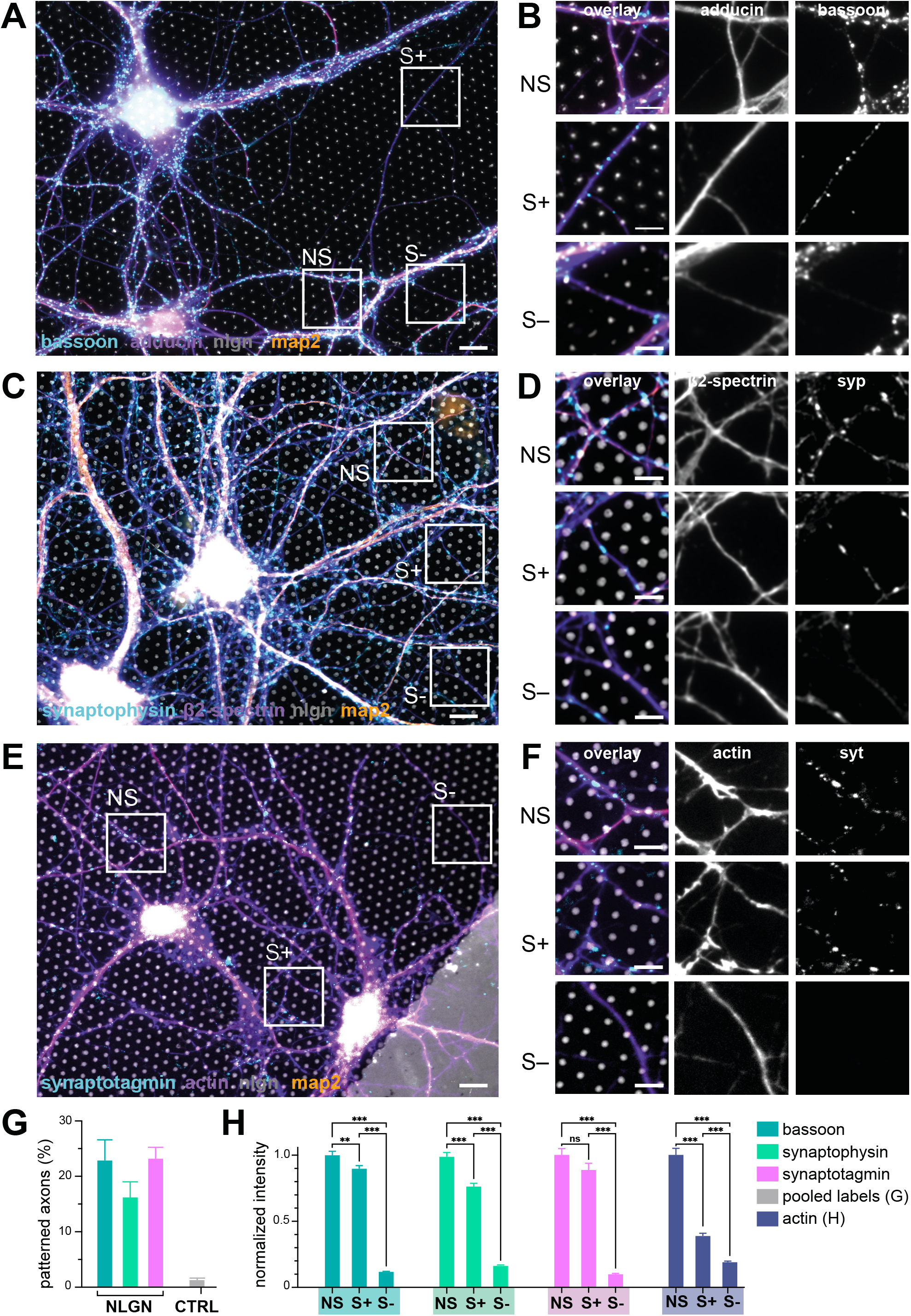
Micropatterned neuroligin induces axonal presynaptic marker clustering and actin aggregation. **A**. Widefield fluorescence image of cultured neurons on top of the nlgn micropattern (gray) fixed at 15 div, labeled for adducin (purple), bassoon (cyan) and map2 (orange). **B**. Zooms corresponding to the areas highlighted in A. Top row: natural synapses (NS) at axo-dendritic contacts; middle row: induced presynapses at isolated axon-micropattern dots (S+); bottom row: isolated axon with no induced presynapse (S-). **C**. Widefield fluorescence image of cultured neurons on top of the nlgn micropattern (gray) fixed at 15 div, labeled for β2-spectrin (purple), synaptophysin (cyan), and map2 (orange). **D**. Zooms corresponding to the NS, S+ and S-highlighted in C. **E**. Widefield fluorescence image of cultured neurons on top of the nlgn micropattern (gray) fixed at div 15, labeled for actin (purple), feeding with anti-synaptotagmin antibody (syt) during constitutive vesicular cycling (cyan) and map2 (orange). **F**. Zooms corresponding to the NS, S+, and S-highlighted in E. Scale bars for A, C, E: 20 µm. Scale bars for B, D, F: 10 µm. **G**. Proportion of the length of isolated axons showing presynaptic specializations following the nlgn micropattern (patterned axon) over the total length of isolated axons selected within the experimental conditions (bassoon, synaptophysin, synaptotagmin) compared to controls (pooled labels, see Fig. S1). **H**. Mean intensity for bassoon (dark cyan, N=2; n>100), synaptophysin (light green, N=4; n>200), synaptotagmin (pink; N=5, n>200) and actin (blue) at natural synapses (NS), induced presynapses (S+), and isolated axon devoid of presynapse (S-), normalized to natural synapses.

On zoomed views, the morphology of bassoon (Fig. 1B), synaptophysin (Fig. 1D), and synaptotagmin uptake (Fig. 1F) clusters are similar between nlgn dot-induced presynapses (noted S+) and natural synapses (noted NS) between axon and dendrites of the same culture. We quantified the signal intensity of each presynaptic marker at ngln dot-induced pre-synapses and compared it to natural synapses and neighboring axonal segments devoid of presynaptic clusters (noted S-, Fig. 1H). Induced presynapses (S+) had a lower intensity compared to NS for bassoon (0.89 ± 0.02) and synaptophysin (0.77 ± 0.02), and the synaptotagmin intensity was not significantly different (0.88 ± 0.04). Control axonal segment (S-) showed very low values of presynaptic component intensity (Fig. 1H). We then measured the actin content of induced presynapses: as expected, the high concentration of postsynaptic actin resulted in an actin intensity being almost 3 times lower in induced presynapses S+ normalized to natural synapses NS (0.38 ± 0.02), but two times higher than in non-synaptic axonal segments S- (0.19 ± 0.01, Fig. 1H), confirming the relative concentration of axonal actin at presynapses (Morales et al., 2000). Altogether, these data show that we can reliably induce the formation of presynapses on micropatterned dots of neuroligin in a minimally perturbed neuronal culture, validating our presynapse-on-glass model.

### Actin enrichment correlates with increased presynaptic component concentration at nlgn dot-induced presynapses

Previous work by our group on bead-induced and natural axon-dendrite presynapses revealed that two categories exist: presynapses showing actin enrichment compared to the surrounding axon (A+) and presynapses lacking actin enrichment (A-). Furthermore, actin enrichment was associated with a higher concentration of presynaptic components and higher cycling activity (Bingham et al., 2023). Consistent with these findings, on micropatterned nlgn dots, induced presynapses could be categorized into actin-enriched and non-enriched from diffraction-limited images (Fig. 2A-B): A+ induced presynapses representing 63% of nlgn dot-induced presynapses, and A-representing 37% (Fig. 2C). We then quantified the enrichment of actin and presynaptic components at nlgn dot-induced presynapses in each category, measuring the fluorescence intensity in A+ and A-induced presynapses as well as the adjacent axonal shaft as a control (S-). Normalized to its intensity in A+ induced presynapses, actin intensity was reduced in A-presynapses and S-shafts (0.38 ± 0.02 and 0.38 ± 0.1, respectively, Fig. 2D). Moreover, A-presynapses demonstrated a significantly reduced concentration of bassoon and synaptotagmin (0.73 ± 0.05 and 0.66 ± 0.05, respectively). Overall, nlgn dot-induced presynapses recapitulate the presence of a majority of actin-enriched presynapses that contain more presynaptic components than their non-enriched counterpart, further validating our presynapse-on-glass model.

**Figure 2:**
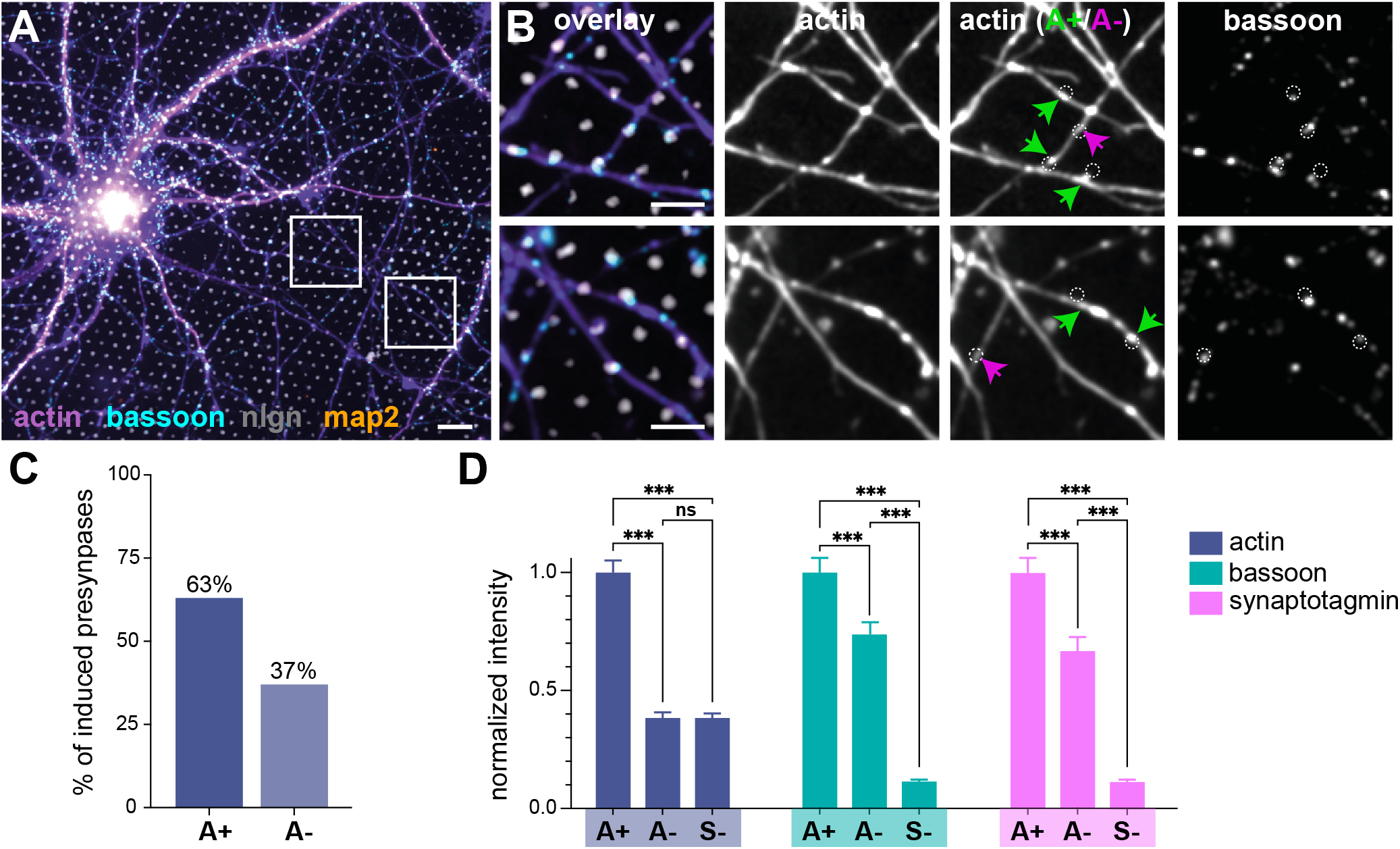
Actin enrichment correlates with increased presynaptic component concentration at nlgn dot-induced presynapses. **A**. Widefield fluorescence image of cultured neurons on top of the nlgn micropattern (gray) fixed at 15 div, labeled for actin (purple), bassoon (cyan), nlgn (gray) and map2 (orange). Scale bars, 20 µm. **B**. Zooms corresponding to the areas highlighted in A displaying (from left to right): overlayed channels; actin; nlgn dots (dashed circles) near actin-enriched induced presynapses (A+, green arrows) and non-enriched presynapses (A-, pink arrows); presynaptic marker bassoon and nlgn dots (dashed circles). Scale bars, 10 µm. **C**. Percentage of actin-enriched and non-enriched induced presynapses along axons on nlgn dots. **D**. Mean intensity of actin (blue; N=3, n>300), bassoon (dark cyan; N=2, n>150), and synaptotagmin (pink, N=3, N>130) at actin-enriched presynapses (A+), non-enriched induced presynapses (A-), isolated axons with no induced presynapses (S-), normalized to actin-enriched presynapses.

### STORM of nlgn dot-induced presynapses reveal discrete actin nanostructures

As noted above, nlgn dot-induced presynapses are visibly distinguishable as A+ and A-at diffraction-limited resolution, particularly when compared side-by-side (Fig. 3A). To further interrogate actin structural organization and assess the presence of the distinct presynaptic actin nanostructures we previously identified in bead-induced presynapses (Bingham et al., 2023), we turned to STORM of actin at nlgn dot-induced presynapses. First, STORM reconstruction confirmed that the periodic actin rings are disrupted at nlgn dot-induced presynapses, as shown previously for the periodic spectrin scaffold at presynapses (He et al., 2016; Sidenstein et al., 2016, Fig. 3B). Furthermore, we were able to identify the three distinct types of actin nanostructures within A+ and A-nlgn dot-induced presynapses (Bingham et al., 2023). Firstly, a large branched and dense structure surrounding the periphery of the presynapse is identified as the actin corral (blue). Secondly, the small linear structures within the bouton often connecting wider and branched structures of the presynapse are classified as actin rails (green). Thirdly, the faint cluster of actin nanostructures colocalizing with the active zone bassoon clusters, is recognized as the actin mesh (Fig. 3B, red). Quantitative analysis of these discrete actin nanostructures showed that all A+ and A-presynapses exhibited a distinguishable peripheral corral. The presence of the fainter rails was detectable in 71% of A+ and 69% of A-presynapses, while the active zone mesh was detected in 83% of A+ and 81% of A-presynapses (Fig. S2). As these numbers were similar for A+ and A-presynapses, we refined our analysis by measuring the area of each nanostructure on STORM images: only the area of corrals showed a significant difference between A+ and A-induced presynapses, with A+ presynapses being larger (0.23 ± 0.01 µm^2^ for A+ vs 0.17 ± 0.03 µm^2^ for A-, Fig. 3C). The presence of all three nanostructures in nlgn dot-induced presynapses, and the main difference between A+ and A-induced presynapses being the relative size of the actin corral surrounding them are perfectly in line with our previous results on bead-induced presynapses (Bingham et al., 2023).

**Figure 3:**
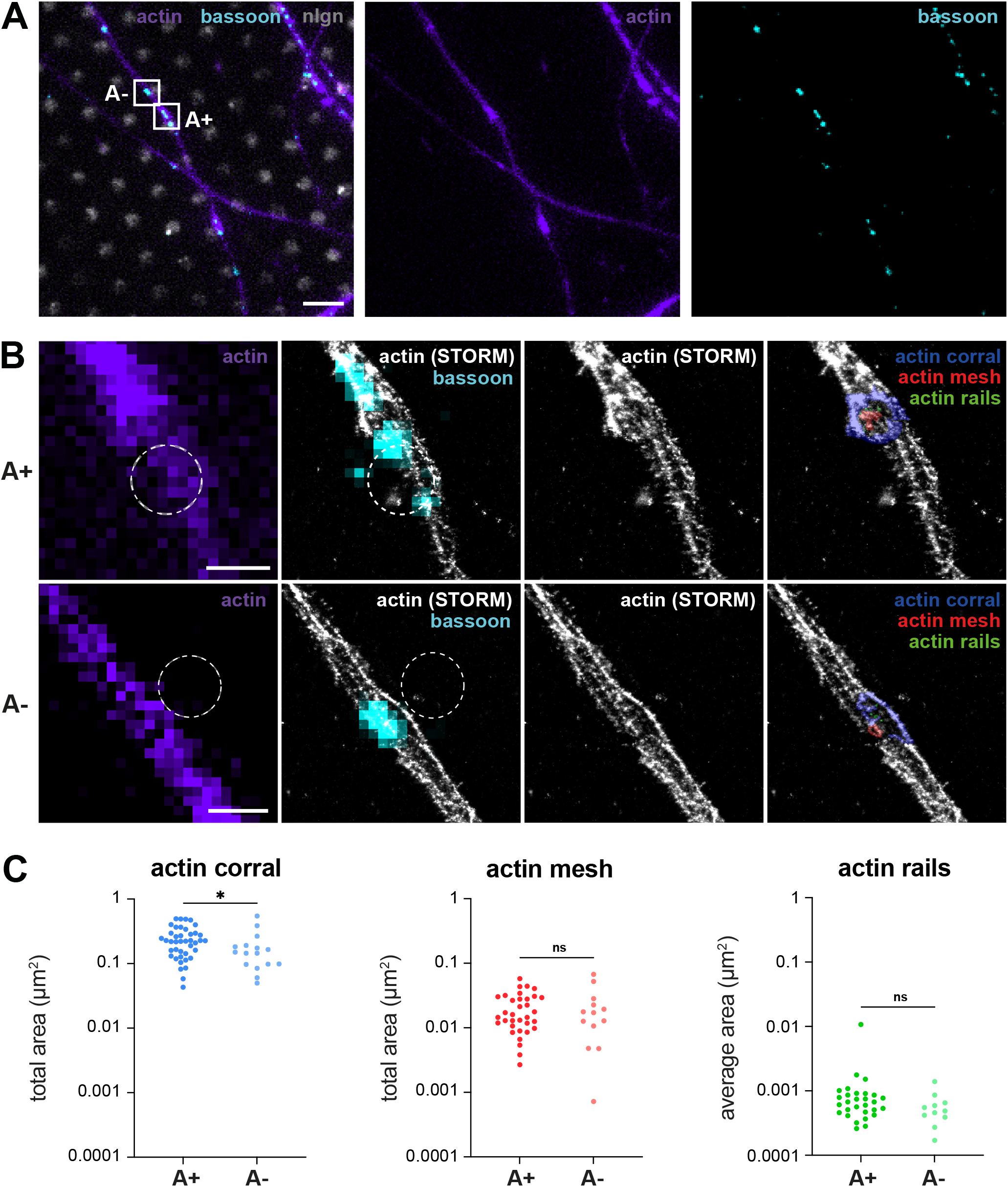
STORM of nlgn dot-induced presynapses reveal discrete actin nanostructures. **A**. Widefield fluorescence image of cultured neurons on top of the nlgn micropattern (gray) fixed at 15 div, labeled for actin (purple) and bassoon (cyan). Scale bar, 4 µm. **B**. Zooms corresponding to two distinct induced presynapses highlighted in A. Left column shows the epifluorescence image of actin (purple) and the position of nlgn dots (white dashed circle). Second column displays the STORM image of actin (gray), epifluorescence bassoon (cyan), and the position of nlgn dots (white dashed circle). Third column shows the STORM image of actin alone (gray). Fourth column shows colorized presynaptic actin nanostructures: corrals (blue), mesh (red) and rails (green). Scale bar, 1 µm. **C**. Areas of presynaptic actin nanostructures as measured on STORM images, comparing A+ and A-induced presynapses. Each point represents an individual measurement, N = 3, n=41 for A+ and 16 for A-.

### Multicolor localization microscopy of nlgn dot-induced presynapses reveals the arrangement of actin nanostructures relative to the active zone

At this point we confirmed that our presynapse-on-glass model recapitulates the results obtained on the bead-induced presynapse model both in terms of actin content and nanostructures. To further show the usefulness of our nlgn dot-induced presynapse, we leveraged the reproducible and controlled orientation of the induced presynapses to interrogate the relationship between actin nanostructures and presynaptic components at the nanoscale. We performed 2-color localization microscopy at nlgn dot-induced presynapses by sequentially combining STORM of actin and DNA-PAINT of the active zone scaffold protein bassoon, using somatodendritic map2 and axonal ß2-spectrin co-staining to target presynapses along isolated axons near nlgn dots (Fig. 4A). We could thus relate the position of the identified actin nanostructures to the bassoon nanoclusters by examining XY and transverse XZ reconstructions (Fig. 4B) as well as ChimeraX renderings (Fig. 4C). Bassoon labeling was made of distinct nanoclusters, as previously described (Glebov et al., 2017). These clusters were located at the bottom of the axon, consistent with the formation of the presynapse toward the glass. The actin corral outlines the periphery of the presynapse, in continuity with rails branching inwardly toward the central part of the presynapse. The structure identified as the mesh based on diffraction-limited views of presynaptic markers (see Fig. 3) could often be seen above the bassoon clusters rather than below them, interdigitating between the clusters to separate them.

**Figure 4:**
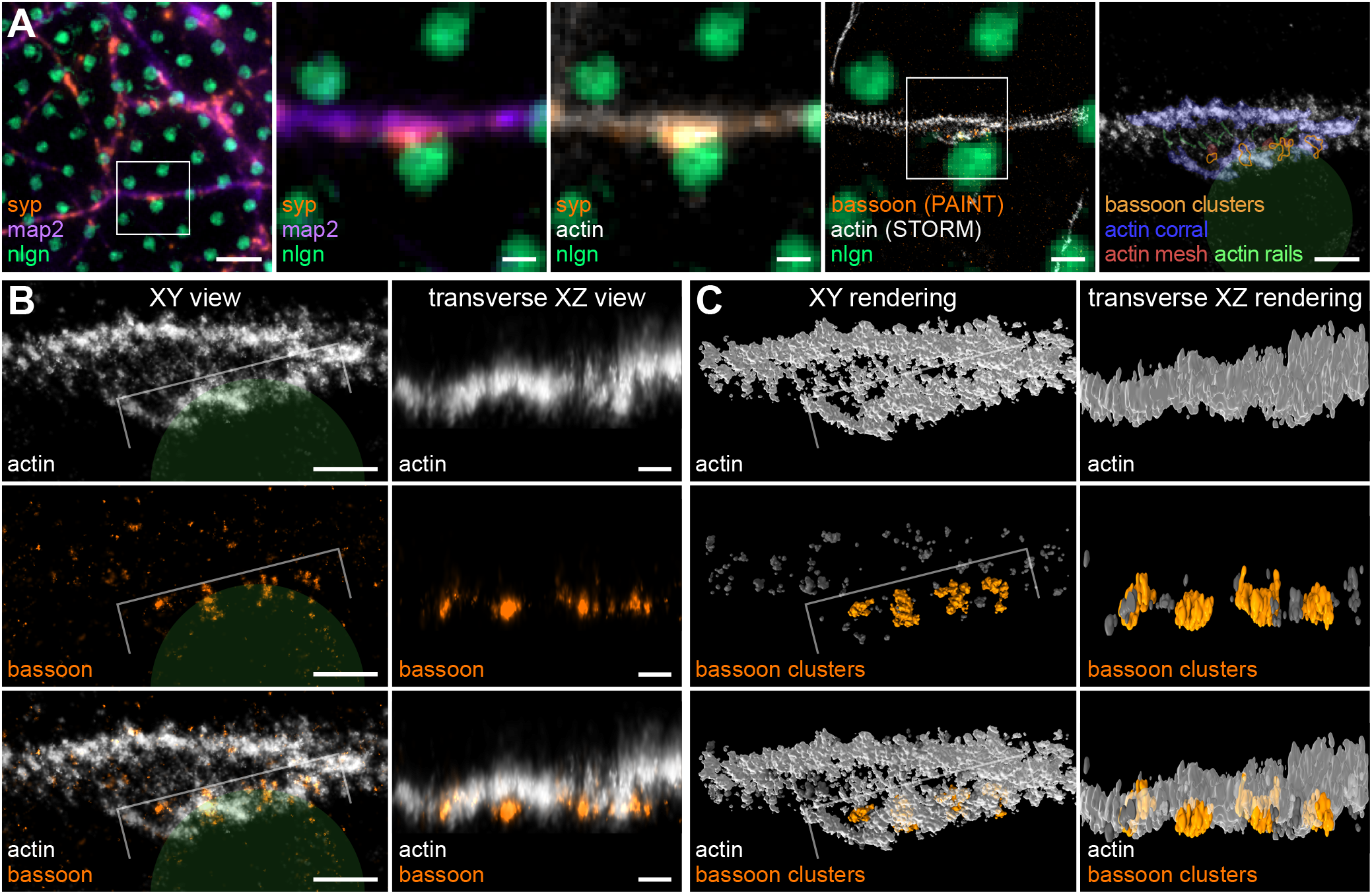
Multicolor localization microscopy of nlgn dot-induced presynapses reveals the arrangement of actin nanostructures relative to the active zone. **A**. First to third images, widefield fluorescence image of cultured neurons on top of the nlgn micropattern (green) fixed at 15 div, labeled for map2 (purple), synaptophysin (orange) and actin (gray). Scale bar, 5 µm (first image), 1 µm (second and third images, zooms of the area highlighted on the first image). Fourth image shows the nlgn pattern (green) superimposed with the PAINT image of the bassoon staining (orange) and the STORM image of the actin staining (gray). Scale bar, 1 µm. Fifth image is a zoom of the area highlighted on the fourth image showing the micropattern (green shape), bassoon clusters from the PAINT image (orange contours), and colorized presynaptic actin nano-structures from the STORM image (gray, corral in blue, mesh in red, rails in green). Scale bar, 500 nm. **B**. Zooms of the presynapse shown in A showing the STORM image of actin (gray) and PAINT image of bassoon (orange) in XY (left column) and transverse (XZ with a vertical Z axis, right column) views. The transverse view is taken along the bracket shown on the XY view. Scale bars, 500 nm (XY views), 200 nm (transverse XZ views). **C**. ChimeraX renderings corresponding to the views shown in B with XY (left column) and transverse (right column) views of actin nanostructures (gray) and bassoon nanoclusters (orange). The transverse XZ views show how actin is present above and in-between bassoon nanoclusters.

### Correlative imaging of the relationship between exocytic sites and presynaptic nanostructures

To further demonstrate the usefulness of our presynapse-on-glass approach, we by took advantage of its controlled *en-face* orientation to set up a live-cell / STORM correlative workflow and resolve the nanoscale environment of synaptic vesicle release events. Neurons were cultured on nlgn-patterned coverslips and transfected with VAMP2-pHluorin, a fusion of the VAMP2/synaptobrevin membrane protein (Baumert et al., 1989) with a pH-sensitive GFP that fluoresces upon exocytosis of the tagged synaptic vesicles (Miesenböck et al., 1998; Burrone et al., 2006), followed by live-cell total internal reflection fluorescence (TIRF) imaging to image spontaneous synaptic vesicle exocytosis at nlgn dots (Fig. 5, first column). The neurons were then fixed and immunolabeled for epifluorescence and single-color STORM imaging of either active zone proteins or presynaptic actin, and the precise location of the exocytic event (from fitting the appearing spot on time-lapse images of VAMP2-pHluorin) was correlated to the nanoscale arrangement from the aligned STORM image (Fig. 5).

**Figure 5:**
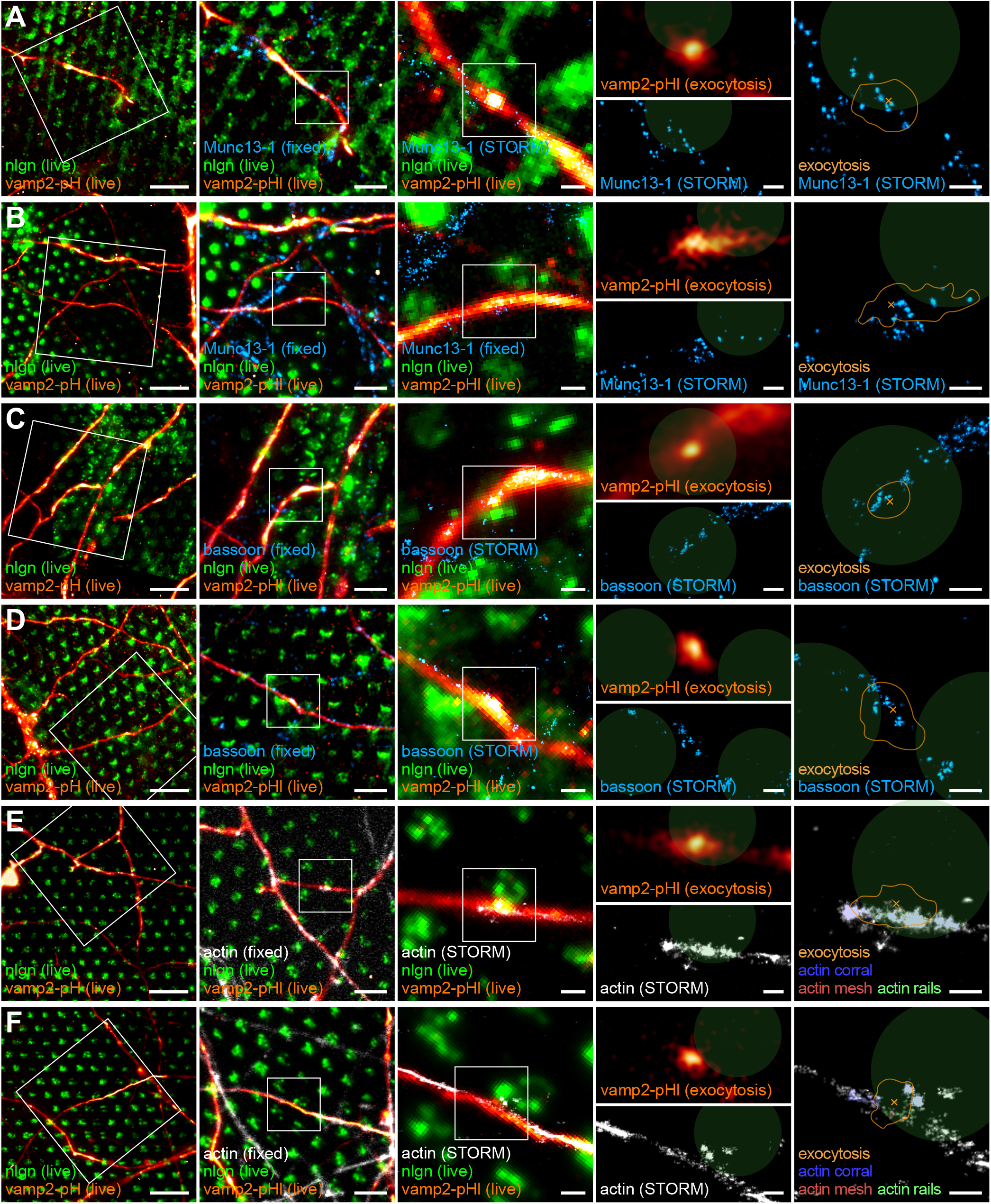
Correlative imaging of the relationship between exocytic sites and presynaptic nanostructures. **A-B**. First image, live-cell imaging of the axon of a cultured neuron at 15 div on top of the nlgn pattern (green), transfected with VAMP2-pHluorin (VAMP2-pHl, orange). Scale bar, 10 µm. Second and third images, progressive zooms on an induced presynapse labeled for Munc13-1 (blue, epifluorescence on the second image, STORM on the third image) after fixation and aligned with the live-cell image. Scale bars, 5 µm (second image), 1 µm (third image). Right panel shows the first frame of the exocytosis event from the VAMP2-pHluorin movie (orange, top left), the corresponding STORM image of the Munc13-1 labeling (blue, bottom left) and the overlay of the exocytosis event (orange contour) with its fitted center (orange cross) with the Munc13-1 STORM image (blue) and nlgn micropattern (green shapes, right). Scale bars, 500 nm. **C-D**. First image, live-cell imaging of the axon of a cultured neuron at 15 div on top of the nlgn pattern (green), transfected with VAMP2-pHluorin (orange). Scale bar, 10 µm. Second and third images, progressive zooms on an induced presynapse labeled for bassoon (blue, epifluorescence on the second image, STORM on the third image) after fixation and aligned with the live-cell image. Scale bars, 5 µm (second image), 1 µm (third image). Right panel shows the first frame of the exocytosis event from the VAMP2-pHluorin movie (orange, top left), the corresponding STORM image of the bassoon labeling (blue, bottom left) and the overlay of the exocytosis event (orange contour) with its fitted center (orange cross) with the bassoon STORM image (blue) and nlgn micropattern (green shapes, right). Scale bars, 500 nm. **E-F**. First image, live-cell imaging of the axon of a cultured neuron at 15 div on top of the nlgn pattern (green), transfected with VAMP2-pHluorin (orange). Scale bar, 10 µm. Second and third images, progressive zooms on an induced presynapse labeled for actin (gray, epifluorescence on the second image, STORM on the third image) after fixation and aligned with the live-cell image. Scale bars, 5 µm (second image), 1 µm (third image). Right panel shows the first frame of the exocytosis event from the VAMP2-pHluorin movie (orange, top left), the corresponding STORM image of the actin labeling (gray, bottom left) and the overlay of the exocytosis event (orange contour) with its fitted center (orange cross) with the actin STORM image (gray) with colored nanostructures (blue, corral; red, mesh; green, rails) and nlgn micropattern (green shapes, right). Scale bars, 500 nm.

As a proof of concept, we first contextualized the exocytic event by localizing presynaptic components with STORM images of Munc13-1, a component of the presynaptic release machinery (Augustin et al., 1999), and the active zone scaffold protein bassoon (Südhof, 2012). Sub-resolution fitting of VAMP2-pHluorin flashes at nlgn-induced presynapses resulted in exocytic localizations in close proximity (<50 nm) to small Munc13-1 clusters (Fig. 5A-B), whereas exocytosis was seen to occur further away from the larger, more uniform bassoon clusters (Fig. 5C-D). We then performed a similar experiment, this time combining live-cell imaging of VAMP2-pHluorin with STORM of actin, counterstaining for bassoon to confirm presynapse induction at individual nlgn dots (Fig. 5E-F). Comparing the localization of single spontaneous exocytic events to labeled actin nanostructures revealed that at the nanoscale, exocytosis occurs in presynaptic regions devoid of actin, presumably emerging from areas adjacent to actin mesh (Fig. E-F). These qualitative results together with the 2-color nanoscale imaging of bassoon and actin (Fig. 4) support an emerging model where actin is interspaced with bassoon clusters to structure the active zone in different nanoscale areas, with further segregation within these areas between bassoon scaffold clusters and Munc13-1/RIM-enriched exocytic zones (Glebov et al., 2017).

## Discussion

In this study, we designed and optimized a presynapse-on-glass model to more precisely investigate actin nano-architecture in induced presynapses. By printing the extracellular domain of neuroligin-1 before coating the coverslips with polylysine and laminin, we developed a micropatterned substrate that promotes natural development of neurons over the whole coverslip. This contrasts with previous methods that coaxed presynapse induction by overexpressing presynaptic neurexin in addition to coating the coverslip with neuroligin (Funahashi et al., 2018; Tanaka et al., 2023) or restrict axon attachment to small spots separated by cell-repellent substrates (Czöndör et al., 2013). Compared to our previous model of presynapses induced by ∼5 µm beads (Bingham et al., 2023), our new model drives presynaptic orientation toward the glass, improving visual accessibility for nanostructure visualization and unbiased exocytosis analysis.

We demonstrate that nlgn dots can induce presynaptic specializations along isolated axons, with concentration of presynaptic components (bassoon, synaptophysin) and active vesicular cycling demonstrated by synaptotagmin antibody uptake. Induced presynapses show a similar morphology, fluorescence intensity of presynaptic components, and vesicular cycling activity compared to natural synapses found at axon-dendrite contacts. The quantification of the fraction of axons exhibiting induced presynapses over nlgn dots might appear low (∼20%), but it is a consequence of our stringent criteria for identifying them, excluding segments with less than three successive induced presynapses to avoid over-estimating induction by including spurious clusters. It should be noted that our micropatterning strategy differs from previously published ones, where the presynapse-inducing patterns (1.5 µm-diameter dots) were complementary with a cytophobic substrate (polyethyleneglycol, Czöndör et al., 2013). In our model, an adhesion-promoting coating of polylysine and laminin is present uniformly, superimposed with the nlgn dots to allow natural growth of neurites over the coverslip with induction of presynapses when axons cross the nlgn dots.

Besides their similarity to natural synapses, the absence of the more intense postsynaptic actin at induced presynapses allowed us to distinguish two populations of presynapses: 63% that are actin-enriched (A+) and 37% that are not enriched (A-), a proportion close to the results obtained on bead-induced presynapses (67% and 33%) and actin-tagged natural presynapses (75% and 25%, Bingham et al., 2023). Furthermore, actin-enriched (A+) induced presynapses similarly accumulated ∼50% more presynaptic components compared to their A-counterparts, validating our nlgn dots-induced presynapse model.

In addition to closely recapitulating the properties of bead-induced presynapses, the presynapse-on-glass model allowed nanoscale visualization with improved optical accessibility due to the formation of an active zone with a controlled orientation and proximity to the substrate. We could thus refine the characterization of the presynaptic actin nanostructures we previously identified (Bingham et al., 2023). Actin corrals are consistently found distributed at the periphery of presynapses, with the corrals of A+ presynapses being significantly larger than those of A-presynapses. These variations in shape and size and the difference in presynaptic content between A+ and A-presynapses suggests that corrals are the primary remodeled actin structure during presynaptic plasticity (O’Neil et al., 2021). Interestingly, we observed a greater percentage of both actin rails and actin mesh in nlgn dot-induced presynapses compared to the bead-induced presynapses (Bingham et al., 2023). This indicates that nlgn dots-induced presynapses indeed exhibit enhanced visibility and allow to more effectively distinguish between the corrals, rails and mesh nanostructures.

One of the key advantages of the presynapse-on-glass model is the controlled orientation of the induced presynapses that form *en face* to the glass substrate. This allows unambiguous interpretation of the 3D nanoscale architecture, without the need for a postsynaptic marker that is often used to determine synapse orientation (Dani et al., 2010; Tang et al., 2016). We could verify the orientation of the induced presynapses by imaging the nanoscale arrangement of bassoon together with presynaptic actin architecture: bassoon nanoclusters were consistently found at the bottom of the axon, even when the presynapse formed on one side of the shaft, and this allowed easier identification of actin corrals, rails and mesh within presynapses. Oriented 3D, multicolor SMLM images allowed us to refine our previous results by demonstrating that the actin mesh, previously coated at the active zone (Bingham et al., 2023) is found above and between bassoon clusters rather than below them. This suggests that rather than a submembrane assembly directly shaping the vesicle exocytosis area, the mesh could have a role in organizing nanoclusters within the active zone, in line with the declustering of bassoon previously observed upon latrunculin A treatment (Glebov et al., 2017).

Being able to visualize synaptic vesicle exocytosis in living cells, including rare spontaneous events, and to correlate them with the nano-architecture of presynaptic component is another application we demonstrate of our presynapse-on-glass model. We visualized individual exocytic events that likely correspond to single-vesicle release (Funahashi et al., 2018) and fit their sub-diffraction localization. By correlating with fixed-cell STORM images of components within the same presynapse, we confirmed that exocytosis occurs next to or at Munc13-1 clusters (Sakamoto et al., 2018), while scaffold proteins such as bassoon are present further away from the exocytic location, in line with their proposed complementary segregation in distinct nanoclusters within the active zone (Tang et al., 2016; Glebov et al., 2017). Furthermore, we used this correlative approach to further understand the role of presynaptic actin. We consistently visualized exocytic events away from actin nanostructures, including the actin mesh, confirming that actin seem to be mostly absent from the exocytic points and further suggesting that active zone actin has a role in defining hotspots of vesicular release where it is absent (Morales et al., 2000). Our results on the nanoscale relationship between actin structures and presynaptic components and vesicular release are primarily qualitative at this point, due to the low throughput of our current correlative approach, and will undoubtedly guide future work by helping to refine functional hypotheses about the roles of actin in presynapses.

## Acknowledgments

We would like to acknowledge funding by the Fédération pour la Recherche Médicale (FRM Equipe grant EQU202103012966) and Agence Nationale pour la Recherche (ANR grant ANR-20-CE13-0024) to C.L. We would like to thank the Neuro-Cellular Imaging Service and Nikon Center for Neuro-NanoImaging at INP, with funding from CPER-FEDER (PlateForme NeuroTimone PA0014842), the Institut Marseille Imaging and NeuroMarseille for complementary equipment funding from Excellence Initiative of Aix-Marseille University – A*MIDEX, a French “Investissements d’Avenir” program (AMX-19-IET-002). We would like to thank Maxime Cazorla (Institut des Neurosciences de la Timone, INT, Marseille, France) for helping with the use of the plasma cleaner, as well as Stephanie Gupton (UNC, Chapell Hill, USA) and Tomoo Hirano (Department of Biophys-ics, Kyoto University, Japan) for the gift of plasmids.

## Author Contributions

Methodology: S.T., L.M., T.W., B.V., M.T., M.-J.P., C.L.; Formal analysis: S.T., L.M., M.-J.P., C.L.; Investigation: S.T., L.M.; Resources: B.V., M.T.; Writing-Original Draft: S.T., L.M., M.-J.P., C.L.; Writing-Review and Editing: S.T., L.M., T.W., M.T., M.-J.P., C.L.; Visualization: S.T., L.M., C.L.; Conceptualization: M.-J.P, C.L.; Supervision, Project Administration: M.-J.P., C.L.; Funding acquisition: C.L.

## Methods

### Plasmids

The expression vector for nlgn (gift from T. Hirano, Funahashi et al., 2018) consists of the cDNA encoding the extracellular region (amino acids 1–675) of neuroligin 1 (−A+B) fused to the N-terminal of human immunoglobulin-Fc region in the pCAGplayII expression vector (Kawaguchi and Hirano, 2006). VAMP2-pHluorin (gift from S Gupton, Urbina et al., 2018) consists in the human VAMP2 fused to super-ecliptic pHluorin (Miesenböck et al., 1998).

### Fc-neuroligin (nlgn) production

Nlgn production followed published procedures (Tanaka et al., 2014). HEK 293 T cells cultured at 70% confluence in 100 mm diameter dishes were transfected with a mixture of Lipofectamine 3000 (Thermo Fisher Scientific; 16 µL/dish) and nlgn plasmid (8 µg/dish) for 4 h in DMEM. Protein expression was carried out for 2 days in DMEM, 10% ultra-low IgG fetal bovine serum (Thermo Fisher Scientific), 100 UI/mL penicillin/streptomycin. The culture medium was then recovered and centrifuged for 10 min at 3000 g to remove cell debris. 200 µL of protein-A Sepharose beads (Sigma) was added to the 50 mL of supernatant in the presence of anti-protease (Sigma) and incubated overnight at 4°C with stirring. After centrifugation for 5 min at 1200 rpm, the beads were washed three times with phosphate buffer saline (PBS, pH 7.4) and nlgn was eluted with 600 µL of 0.1M glycine buffer, pH 2.6 and immediately buffered to pH 7 with 1M Tris, pH 9. Purified nlgn concentration determined by dot blot was consistently around 250 µg/mL.

### Micropatterned substrate onto glass coverslips

18-mm diameter round, #1.5H thickness coverslips were rapidly washed in acetone and ethanol, then affixed with dots made from two-components orthodontics paste (Rotec) as spacers. They were then oxidized in a plasma cleaner (Fento, Diener Electronic). Micro-contact printing was performed using PDMS stamps with 1.8 µm-diameter pillar arrays with 4 µm center-to-center distance (gift of Jianping Fu, University of Michigan, USA, Weng and Fu, 2011) according to a previously published protocol (Théry and Piel, 2009). Stamps were coated and incubated for 40 min at 37°C with a mixture of BSA-Alexa Fluor 488 (80 µg/mL; Thermo Fisher Scientific) and nlgn (25 µg/mL), or with BSA-AF488 (80 µg/mL) mixed with the Fc fragment of human IgG (25 µg/mL; Jackson ImmunoResearch) as a control. PDMS stamps were then inverted onto ionized coverslips and incubated at 37°C for 20 min, then treated overnight with a mixture of laminin (Thermo Fisher Scientific; 10 µg/mL) and poly-L-lysine (Sigma; 38 µg/mL), prior to neuron seeding.

### Animals and cell culture

The use of Wistar rats followed the guidelines established by the European Animal Care and Use Committee (86/609/CEE) and was approved by the local ethics committee (agreement G13055). Rat hippocampal neurons were cultured following the Banker method, above a feeder glia layer (Kaech and Banker, 2006). Hippocampi from E18 rat pups were dissected and homogenized by trypsin treatment followed by mechanical trituration and seeded on the patterned coverslips at a density of 4,000-6,000 cells/cm^2^ for 3 h in serum-containing plating medium (MEM with 10% FBS, 0.6% glucose, 0.08 mg/mL sodium pyruvate, and 100 UI/mL penicillin/streptomycin). Coverslips were then transferred, cells down, to petri dishes containing confluent glia cultures conditioned in neurobasal medium (NB; Thermo Fisher Scientific) supplemented with 2% B27 (Thermo Fisher Scientific), 100 UI/mL penicillin/streptomycin, and 2.5 µg/mL amphotericin, and cultured in these dishes at 37°C, 5% CO_2_. Neurons were fixed at 13-16 days in vitro (div).

### Anti-synaptotagmin vesicular cycling assays

Vesicular cycling assays were performed using an antibody directed to an extracellular epitope of synaptotagmin 1 (syt, clone 604.2; Synaptic Systems, Kraszewski et al., 1995) that was fed to neurons before fixation. Neurons were incubated for 1 h at 37°C, 5% CO_2_ with the syt antibody at 1.6 µg/μL in HBS medium (136 mM NaCl, 10 mM HEPES, 10 mM glucose, 2 mM CaCl_2_, 1.3 mM MgCl_2_). Cells were then washed three times with PBS before fixation.

### Antibodies

Primary antibodies used are chicken anti-map2 (1:1,000, ab5392, RRID:AB_2138153; Abcam), mouse anti-ß2-spectrin (1:100, clone 42/aa 2101-2189, #612563, RRID:AB_399854; BD Bioscience), guinea pig anti-synaptophysin (1:400, 101 004, RRID:AB_1210382; Synaptic Systems), mouse anti-synaptotagmin1 (1:600, clone 604.2/aa 1-12, 105 311, RRID:AB_993036; Synaptic Systems), rabbit anti-adducin alpha (1:100, ab40760, RRID:AB_722627; Abcam), mouse anti-bassoon (1:200, clone SAP7F407, ab82958, RRID:AB_1860018; Abcam), rabbit anti-GFP (1:750, ab290, RRID:AB_303395; Abcam), mouse anti-GFP (1:750, clone 9F9.F9, ab1218, RRID:AB_298911; Abcam), rabbit anti-munc13-1 (1:500, 126 103, RRID:AB_887733; Synaptic Systems). Secondary antibodies used were conjugated to Alexa Fluor 488, 555, or 647 (Thermo Fisher Scientific), Atto 643 (Massive Photonics), or DyLight 405 (Rockland) and diluted 1:200-1:400. Actin was labeled with phalloidin Alexa Fluor 647 Plus (1:400, Thermo Fisher Scientific).

### Fluorescence immunocytochemistry

Immunocytochemistry of neuronal cultures for widefield microscopy, SMLM and live-cell imaging was performed as in published protocols (Jimenez et al., 2020) with minor modifications. Cells were fixed using 4% PFA in PEM buffer (80 mM PIPES pH 6.8, 5 mM EGTA, 2 mM MgCl_2_) for 10 min at room temperature (RT). After rinses in 0.1 M phosphate buffer (PB), neurons were blocked for 1-2h at RT in immunocytochemistry buffer (ICC: 0.22% gelatin, 0.1% Triton X-100 in PB), and incubated with primary antibodies diluted in ICC for either 1h30 at RT or overnight at 4°C. After rinses in ICC, neurons were incubated with secondary antibodies diluted in ICC for 1h at RT and rinsed. Actin staining was then performed by incubating in fluorescent phalloidin at 0.5 μM for either 1h at RT or over-night at 4°C. Stained coverslips were kept in PB + 0.02% NaN_3_ at 4°C until imaging. For widefield microscopy, coverslips were mounted in ProLong Glass (Thermo Fisher Scientific).

### Epifluorescence microscopy

Diffraction-limited epifluorescence images were obtained using an Axio-Observer upright microscope (Zeiss) equipped with a 40X NA 1.4 or 63X NA 1.4 objective and an Orca-Flash 4.0 camera (Hamamatsu). Appropriate hard-coated filters and dichroic mirrors were used for each fluorophore. A thin Z-stack of 3-10 slices spaced by 0.2 µm was acquired to include the signal from all neuronal processes within the whole field of view. For illustration images, image editing was performed using Fiji (Schindelin et al., 2012) with linear contrast adjustment to highlight the faint actin labeling along axons.

### Epifluorescence image analysis

Percentage of patterned axon quantifications were performed by defining linear regions of interest (ROIs) and measuring their length using the Fiji software. Patterned axon ROIs were identified along isolated axons (devoid of dendritic contact) as presenting at least three successive presynaptic aggregates next to nlgn dots; other isolated axon segments were defined as non-patterned. The percentage of patterned axons was calculated over the total length (patterned + non-patterned ROIs).

Intensity quantifications were performed on maximum projections of the raw epifluorescence data with no further adjustment. Short linear ROIs were defined onto the presynaptic cluster aggregates at nlgn dot-induced presynapses (S+), at natural (axon-dendrite) synapses (NS), and at the axonal shaft flanking the presynapses as control (S-). Those tracings were translated into Fiji ROIs and refined to the presynaptic cluster (S+ and NS) or axon shaft (S-) using the ProFitFeat script (available at https://github.com/cleterrier/Measure_ROIs/blob/master/Pro_Feat_Fit.js), limiting it to the segment with an intensity above 40% of the maximum intensity along the linear ROI. The background-corrected mean intensity along these ROIs was then measured for each labeled channel. Induced presynapses were visually categorized as “actin enriched” (A+) or “non-enriched” (A−) depending on the relative intensity of actin compared to the neighboring axon shaft (Bingham et al., 2023).

### SMLM: STORM and PAINT

Both STORM and DNA-PAINT acquisitions were performed on an N-STORM microscope (Nikon Instruments). The N-STORM system uses an Agilent MLC-400B laser launch with 405 nm (50 mW maximum fiber output power), 488 nm (80 mW), 561 mW (80 mW), and 647 nm (125 mW) solid-state lasers, a 100X NA 1.49 objective, and an Ixon DU-897 camera (Andor). After locating a patterned axon using low-intensity epifluorescence illumination, a STORM or DNA-PAINT acquisition was performed using laser illumination in HiLo (grazing angle) configuration. An astigmatic lens was added to the light path to achieve 3D imaging. For STORM, stained coverslips were mounted in a silicone chamber filled with STORM buffer (Smart Buffer Kit, Abbelight), and 30,000-60,000 images (256 × 256 pixels, 15 ms exposure time) were acquired at 100% 647-nm laser power. Reactivation of fluorophores was performed during acquisition by increasing illumination with the 405-nm laser. For DNA-PAINT, stained coverslips were mounted in a Ludin chamber filled with an imaging buffer (Massive Photonics). Imaging strands conjugated to Atto643 (Massive Photonics) were added from a starting concentration of 0.1 nM each, then adjusted to optimize blinking density. 30,000-45,000 images (256×256, 40 ms exposure time) were acquired at 50-60% laser power for 647 nm laser excitation.

To obtain optimal STORM images of presynaptic actin, we carefully selected induced presynapses with unambiguous proximity with a nlgn dot along an isolated, continuous axon. We avoided situations where the nlgn dot was partially printed or erased, as well as regions of axons crossing or bundling.

For STORM and DNA-PAINT images, acquired stacks were processed using DECODE (Speiser et al., 2021). Briefly, PSFs were modeled using spline fitting in SMAP (Li et al., 2018b) and used to simulate sequences of blinking events using characteristics (photon number range and lifetime distribution) inferred from real acquisition data from the N-STORM microscope. A PyTorch model was trained to infer the 3D coordinates and uncertainty of the simulated blinking events and then applied to the experimental acquired sequence. The resulting localizations (fitted blinking events) were filtered based on uncertainty, and drift during acquisition was corrected in 3D using a redundant cross-correlation algorithm (Wang et al., 2014) implemented as an independent module of SMAP (Ries, 2020). After translation of the coordinate files, image reconstructions were performed using the ThunderSTORM ImageJ plugin (Ovesný et al., 2014) in Fiji software. Custom scripts and macros were used to translate coordinate files and automate image reconstruction for images at 16 nm/pixel for visualization and at 8 nm/pixel for cluster analysis (https://github.com/cleterrier/ChriSTORM).

### STORM image analysis

A high-magnification image (4 nm/pixel) was generated for each induced presynapse image, and manually registered by translation with the corresponding epifluorescence images in other channels, including presynaptic marker. Actin structures present at the induced presynapse were categorized into three types using the following criteria (Bingham et al., 2023):

- actin mesh: a small cluster of actin next to the nlgn dot contact and within the presynaptic active zone marker cluster.
- actin rails: linear structures within the presynaptic marker cluster
- actin corrals: large actin clusters at the periphery or just surrounding the presynaptic marker cluster.

The size of the actin nanostructures was determined using manual outlines on 2D projection of actin STORM images in Fiji, aided by visualization of 3D renderings using ChimeraX software (Goddard et al., 2018).

### Neuronal transfection and live-cell imaging

13 div cultured neurons, at a density of 6,000 cells/cm^2^, were transfected using Lipofectamine 2000 (Thermo Fisher Scientific) with 0.15 μg of VAMP2–pHluorin. Following 30 min incubation in neurobasal medium (NB; Thermo Fisher Scientific) at 37oC, 5% CO_2_, coverslips were returned to their original culture dishes. After 16 h, live-cell imaging of neurons expressing VAMP2–pHluorin was performed on an inverted microscope ECLIPSE Ti2-E (Nikon Instruments) equipped with an OR-CA-Fusion sCMOS camera (Hamamatsu Photonics) and a CFI SR HP Apochromat TIRF 100× oil (NA 1.49) objective. The system was equipped with a Nikon Perfect Focus System (PFS) and images were acquired with the NIS-Elements software. Coverslips with neurons were mounted in a Ludin metal chamber in Tyrode’s solution (119 mM NaCl, 25 mM HEPES, 2.5 mM KCl, 2 mM CaCl_2_, 2 mM MgCl_2_, 30 mM glucose, pH 7.35). Neurons were maintained in a humid chamber at 35.5–37°C for the duration of the experiments using a stage-top incubator (Okolab Inc). To image exocytosis, VAMP2–pHluorin initially present at the plasma membrane was photobleached using high power 488 nm laser light for 15 s before time-lapse imaging to isolate exocytic signal (Yudowski et al., 2007); 100 ms exposure frames were then continuously acquired for 5 min using 488 nm laser light at lower power (1–10%).

### Timelapse data processing

Acquired videos were preprocessed using the Filter Timelapse script (available at https://sites.imagej.net/Christopheleterrier/plugins/NeuroCyto%20Lab/Kymographs/), which performed image stabilization using the Image Stabilizer plugin (https://imagej.net/plugins/image-stabilizer) after 2× downscaling, and bleach correction using average intensity compensation for the foreground identified as the top 12% of pixel intensities within each image of the sequence. Exocytosis events along isolated axons across a field of nlgn dots were then identified with the pHusion plugin in ImageTank software (O’Shaughnessy et al., 2024). Exocytotic events were detected by using the Difference of Gaussian (DoG) method using a sigma of 3 and a scaling factor of 1.25. Using the region of interest from the DoG image, a small image series was cropped from the raw image stack taking 5 frames before and 30 after the identified spot. Each image in the series was fitted with a gaussian model. Only events with a goodness of fit for the gaussian model greater than 0.7 were considered. Events that drifted greater than 4 pixels were removed. Identified exocytosis events were subsequently verified by visual inspection of the original timelapse movies in Fiji.

### Correlative live-cell/STORM

Live-cell timelapse data and corresponding STORM images were registered with a two-step procedure. First, an RGB merge of the maximum-projected VAMP2-pHluorin movie and nlgn pattern from live-cell data was aligned to an RGB merge of the fixed VAMP2-pHluorin image from the STORM acquisition using a manually-tuned affine transform using the BigDataViewer Fiji plugin (Pietzsch et al., 2015). The resulting affine transform was then applied to the live-cell data using the TransformJ Fiji plugin to align it to STORM reconstructions and corresponding up-sampled epifluorescence channels (at 16 or 4 nm per pixel depending on the zoom level). Vamp2-pHluorin timelapse was up-sampled using a quintic B-spline approach (Meijering et al., 2001), and the exocytic localization was determined on the first frame where the exocytosis signal appeared, as the highest intensity pixel on the image up-sampled to 16 nm per pixel.

### Statistics

Individual measurement points (n) from independent experiments (N) were pooled. Intensity profiles, graphs, and statistical analyses were generated using Prism. On bar graphs, dots (if present) are averages of each independent experiment, bars or horizontal lines represent the mean, and vertical lines are the SEM unless otherwise specified. Significances were tested using one-way, non-parametric ANOVA with Šídák post-hoc significance testing between selected conditions (Fig. 1 and 2) or using Wilcoxon rank sum test (Fig. 3). On Figures, the results of the post-hoc significance are indicated as follows: ns or ns, non-significant; *, p < 0.05; **, p < 0.01; ***, p < 0.001.

## Supplementary material

**Figure S1:**
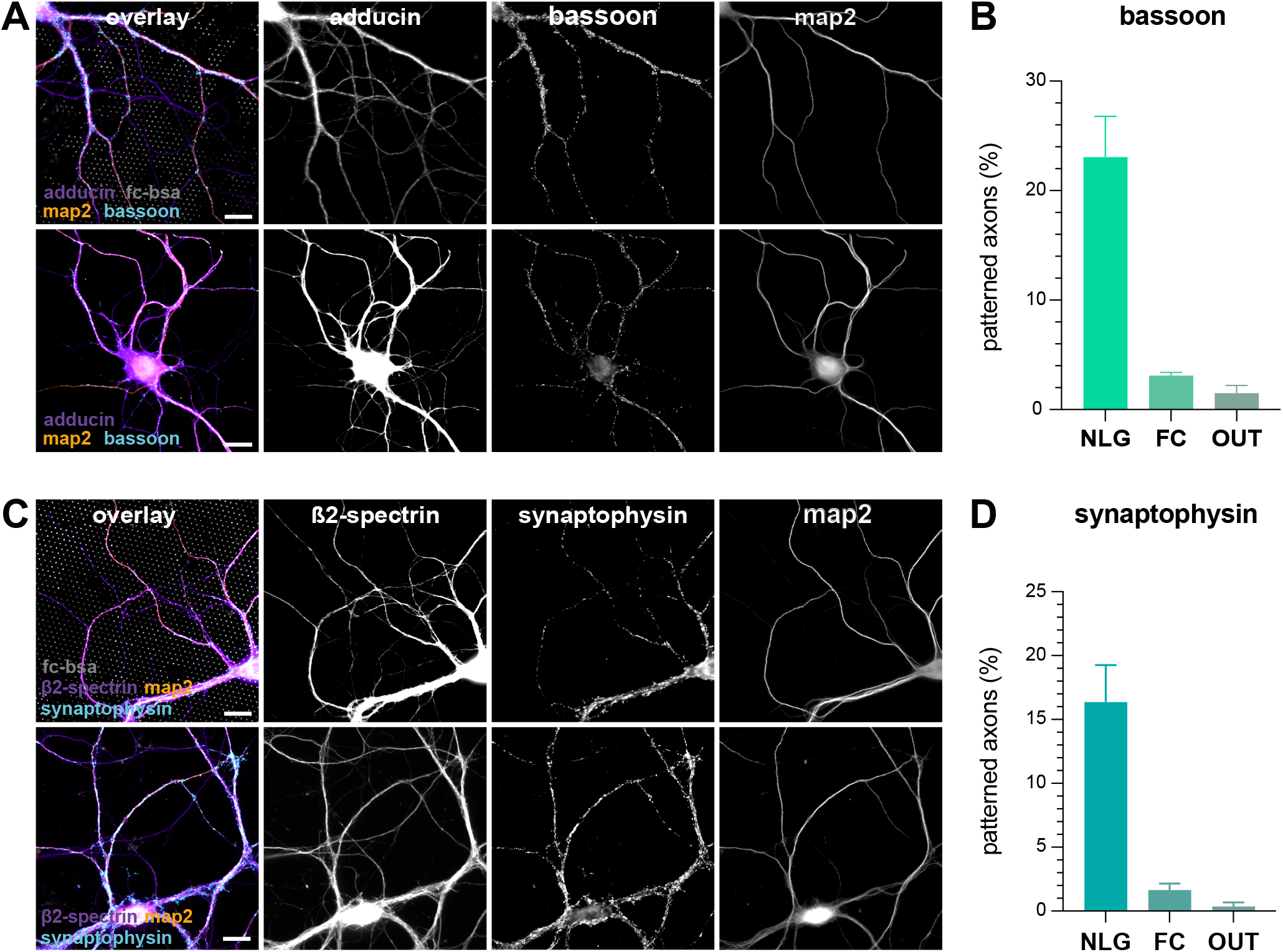
No induction of presynapses on Fc-BSA dots or outside the nlgn dots pattern. **A**. Widefield fluorescence image of cultured neurons on top of the Fc-BSA micropattern (top row, gray) or outside of the pattern (bottom row) fixed at div 15 labeled for adducin (purple), bassoon (cyan), map2 (orange). Scale bar, 10 µm. **B**. Proportion of the length of isolated axons showing presynaptic bassoon specializations following the nlgn micropattern (patterned axon) over the total length of isolated axons selected within the experimental conditions (nlgn dots, N=5) compared to controls (Fc-BSA, N=3; outside of the nlgn dots pattern, N=4). **C**. Widefield fluorescence image of cultured neurons on top of the Fc-BSA micropattern (top row, gray) or outside of the pattern (bottom row) fixed at div 15 labeled for ß2-spectrin (purple), synaptophysin (cyan), map2 (orange). Scale bar, 10 µm. **D**. Proportion of the length of isolated axons showing presynaptic synaptophysin specializations following the nlgn micropattern (patterned axon) over the total length of isolated axons selected within the experimental conditions (nlgn dots, N=4) compared to controls (Fc-BSA dots, N=3; outside of the nlgn dots pattern, N=4).

**Figure S2:**
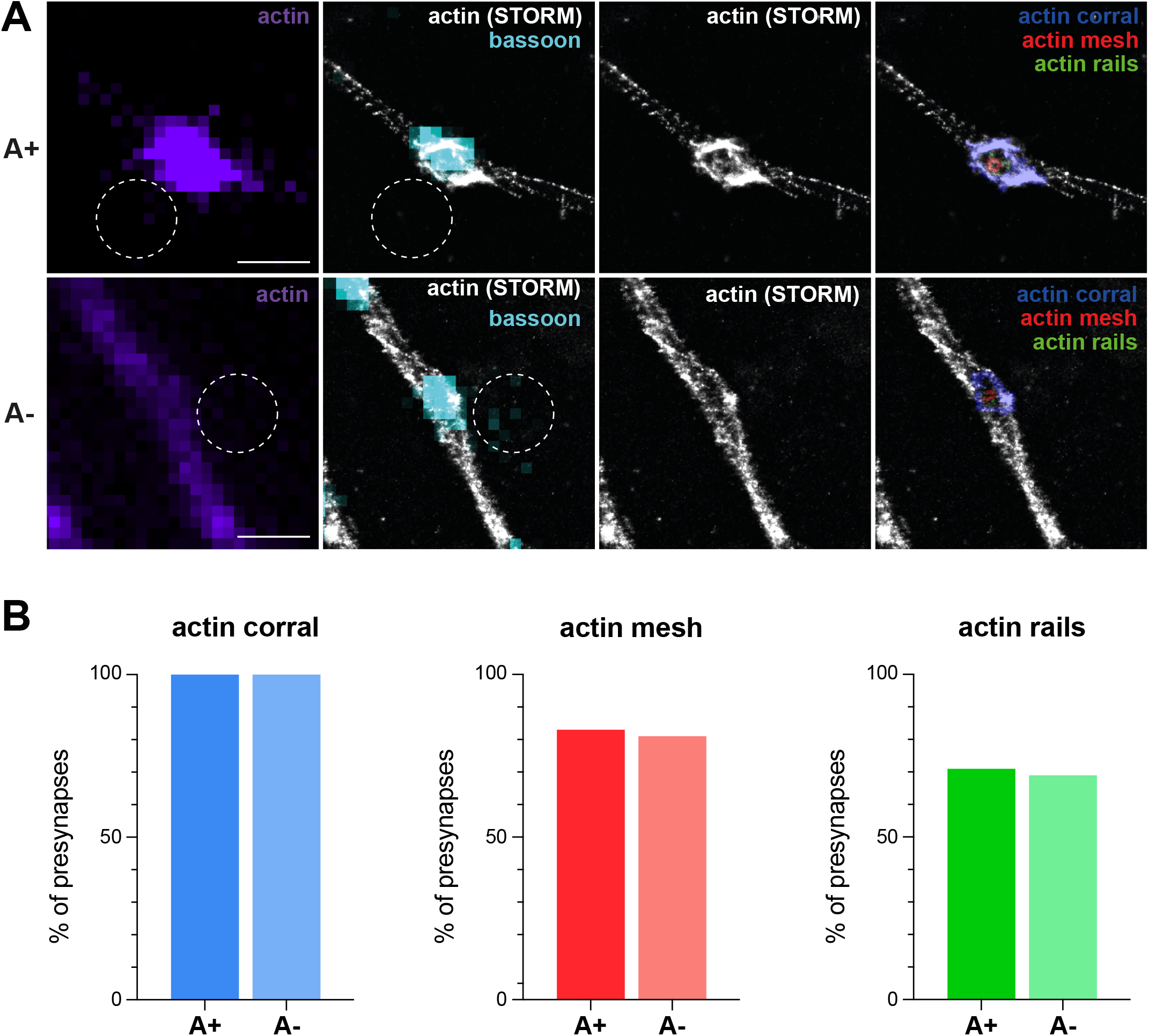
Presence of actin nanostructures in nlgn dot-induced presynapses enriched and non-enriched in actin. **A**. Axon segments displaying actin-enriched (A+, first row) and actin-non-enriched presynapses (A-, second row). The left column displays the epifluorescence image of actin (purple) and the position of the nlgn dots (white dashed circle). Second column displays the STORM image of actin (gray), the epifluorescence image of bassoon (cyan) and the position of the nlgn dots (dotted circle in white). Third column shows the STORM image of actin (gray) alone. Fourth column shows colorized presynaptic actin nanostructures: actin corrals (blue), actin mesh (red) and actin rails (green). Scale bars, 1 µm. **B**. Proportion of presynapses where each nanostructure is detected for A+ and A-induced presynapses: actin corrals (blue), actin mesh (red), and actin rails (green).

